# Mechanism-based approach in designing patient-specific combination therapies for nonsense mutation diseases

**DOI:** 10.1101/2024.11.13.623453

**Authors:** Saleem Bhat, Arpan Bhattacharya, Hong Li, Xianon Cui, John D. Lueck, Yale E. Goldman, Barry S. Cooperman

## Abstract

Premature termination codon (PTC) diseases, arising as a consequence of nonsense mutations in a patient’s DNA, account for approximately 12% of all human disease mutations. Currently there are no FDA approved treatments for increasing PTC readthrough in nonsense mutation diseases, although one translational readthrough inducing drug, ataluren, has had conditional approval for treatment of Duchenne muscular dystrophy in Europe and elsewhere for 10 years. Ataluren displays consistent low toxicity in clinical trials for treatment of several different PTC diseases, but its therapeutic effects on such diseases are inconsistent. The identity of the stop codon and its sequence context are major determinants of PTC readthrough efficiency in both the absence and presence of nonsense suppressors. Previously we have shown that ataluren stimulates readthrough exclusively by competitively inhibiting release factor complex (RFC, eRF1.eRF3.GTP)-dependent catalysis of translation termination. Here, using an *in vitro* reconstituted system (PURE-LITE) and both ensemble and single molecule assays, we demonstrate that PTC identity and the immediately adjacent mRNA sequence contexts modulate the catalytic activity of RFC in terminating peptide elongation. Such modulation largely determines the effectiveness of ataluren in stimulating readthrough, whether added alone or in combination with either the aminoglycoside G418 or an anticodon edited aa-tRNA, each of which stimulate readthrough by mechanisms orthogonal to that of ataluren. Our results provide an attractive rationale for the variability of ataluren effectiveness in stimulating readthrough in clinical trials. Patients harboring a PTC mutation with a sequence context promoting strong interaction with RFC are predicted to be resistant to ataluren, whereas ataluren treatment should be more effective for patient sequences conferring weaker interaction with RFC.

## INTRODUCTION

Premature termination codon (PTC) diseases, arising as a consequence of nonsense mutations in a patient’s DNA, account for approximately 12% of all human disease mutations, including those giving rise to cystic fibrosis (CF), Duchenne muscular dystrophy (DMD), Marfan syndrome (MFS) and several cancers ^**1-4**^. Despite their prevalence, there are currently no FDA approved treatments for increasing PTC readthrough in nonsense mutation diseases. The situation is similar worldwide, although since 2014, the European Medicines Agency and several other national regulatory agencies have given conditional approval of the translational readthrough inducing drug (TRID) ataluren (also known as PTC124 and marketed as Translarna) for treatment of DMD, the only such TRID approved for treatment of any PTC disease. Ataluren has been shown to stimulate readthrough of a PTC, resulting in the insertion of one of several different amino acids, depending on the identity of the pathological nonsense codon (UGA: W, R, or C; UAA or UAG: Q, K, or Y)^**5-8**^. Ataluren displays consistent low toxicity in clinical trials for treatment of several different PTC diseases, but its therapeutic effects on such diseases are inconsistent^**9**^.

The lack of treatments for PTC diseases which increase readthrough has spurred research to develop clinically relevant nonsense suppressors^**10-14**^. These include small organic molecule TRIDs, <u>a</u>nti<u>c</u>odon-<u>e</u>dited suppressor tRNAs (ACE-tRNAs), which have recently been shown to promote readthrough of disease-causing PTC mutations to an impressive extent in both cellular and animal studies^**15-20**^, and mRNA and DNA editing^**21,22**^. Another active field of research focuses on how the identity of the stop codon and its sequence context affect readthrough efficiency (RE), in both the absence and presence of nonsense suppressors^**23** - **27**^. It is clear from these studies, mostly in cells or cell extracts, that readthrough is most pronounced with the UGA PTC and that both downstream and upstream sequences significantly affect RE. Recent results of Toledano et al.^**23**^ and Mangkalaphiban et al.^**24, 25**^, provide strong evidence that, in addition to the identity of the PTC itself (UGA, UAA, or UAG), by far the strongest effects of the nearby sequence context depend on one and possibly two codons immediately downstream (nts +4 - +9) and one codon upstream (nts -1 to -3) of the stop codon (nts +1 to +3). The downstream context is the more consequential ^**23-25, 28**^, likely reflecting in part variation in the direct interaction strength of the +4 nt, and possibly the +5 nt, with eukaryotic release factor 1 (eRF1)^**29**^. The upstream codon also plays a substantial role, with both Toledano et al.^**23**^ and Mangkalaphiban et al.^**24, 25**^ concluding that the identity of the P-site peptidyl-tRNA bound to this codon is primarily responsible for upstream effects on readthrough. Lesser effects result from variation in nts -4 to -6.

Our interest in developing potent and clinically useful treatments of PTC diseases has led us to focus on ataluren, because of its low toxicity and its conditional approval for treatment of DMD. Recently, we developed a reconstituted *in vitro* eukaryotic system, denoted PURE-LITE, to conduct detailed mechanistic studies of eukaryotic protein synthesis^**30-32**^. PURE-LITE takes advantage of the ability of the intergenic IRES of Cricket Paralysis Virus (CrPV-IRES) to form a tight complex with 80S ribosomes, which is then capable of initiating cell-free synthesis of complete proteins in the absence of the complex set of natural initiation factors^**33**^. This allows for elongation and termination to be studied with the addition of just four factors, eEF1A and eEF2 for elongation and eRF1 and eRF3 for termination (Figure 1A). Using the PURE-LITE system we recently reported that ataluren stimulates readthrough exclusively by competitively inhibiting release factor complex (RFC, eRF1.eRF3.GTP)-dependent catalysis of translation termination,, and does so via binding to at least two sites on the ribosome and possibly one on the RFC^**32**^. It thus has a mode of action orthogonal to that of the aminoglycoside G418, which, on binding to its high affinity site near the ribosome decoding site, stimulates readthrough by facilitating productive binding of near-cognate suppressor tRNAs (Sup-tRNAs) without affecting termination activity^**31**^ (Figure 1B).

**Figure 1.**
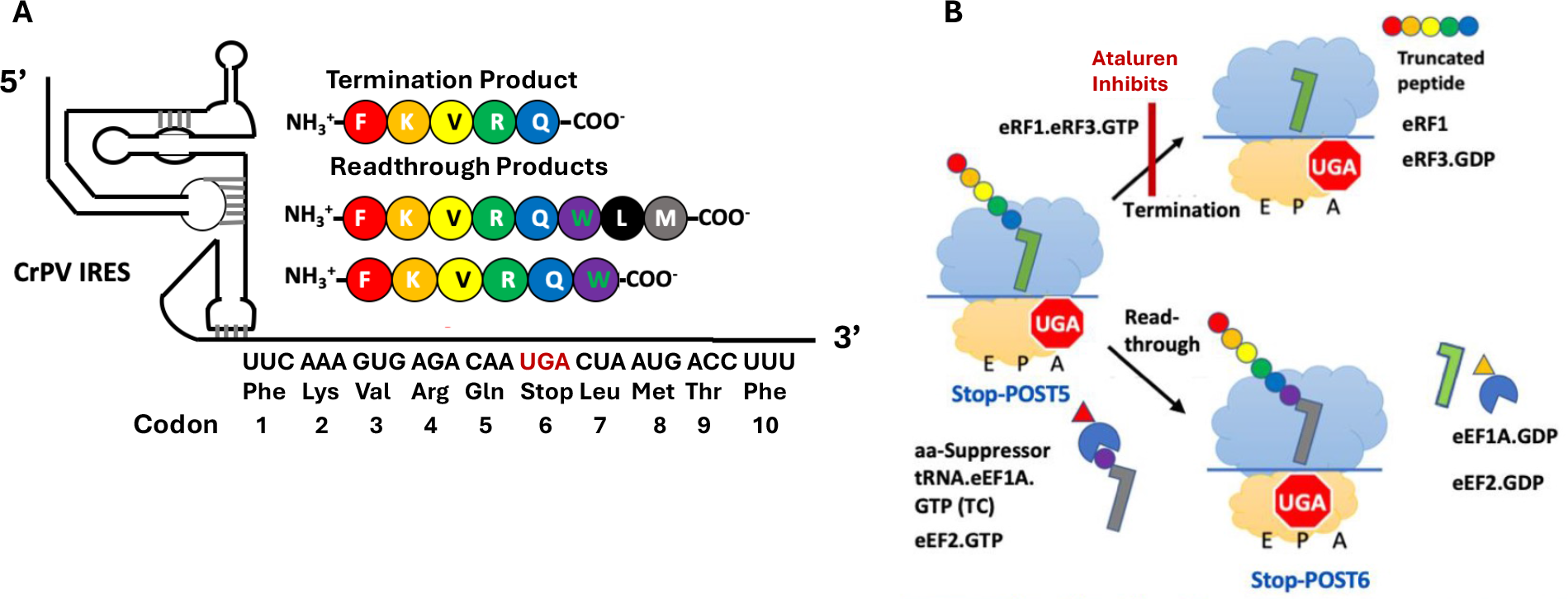
***A***. The Pure-LITE System for Measuring Termination and Readthrough. ***B***. RFC and Suppressor TC compete for reaction with Stop-POST5 pretermination complex. For readthrough, Trp-tRNA_Trp_ is used as suppressor aminoacyl-tRNA.

Here we present results demonstrating that **i**. RFC termination activity has a strong dependence on the sequence context of PTCs found in patient mRNA sequences encoding cystic fibrosis transmembrane conductance regulator (CFTR) and fibrillin1 (FBN1), causing cystic fibrosis and Marfan syndrome, respectively; **ii**. sequence dependent RFC activity correlates significantly with ataluren inhibition of termination and stimulation of readthrough; and **iii**. combinations of ataluren and either of the nonsense suppressors ACE-tRNA^Arg^ and G418 have additive and occasionally synergistic effects on readthrough.

## MATERIALS AND METHODS

### Preparation of ribosomes, elongation factors and aa-tRNAs

Stop-POST5 complexes listed in Tables 1 and 2 and used in termination and readthrough assays were prepared from shrimp (*Artemia salina*) 80S via a high KCl treatment of 80S ribosomes programmed with CrPV-IRES-mRNAs and the ternary complexes (TCs; aa-tRNA.eEF1A.GTP) of the appropriate aminoacyl-tRNAs, as described previously for preparing Reference Stop-POST5.^**32**^. *S. cerevisiae* elongation factors eEF1A^**34**^ and eEF2^**31,35**^ and human eRF1 and eRF3 termination factors were prepared as reported previously^**36,37**^. In the initial phases of this work, yeast tRNA^Phe^ was purchased from Sigma-Aldrich and other isoacceptor tRNAs were prepared from bulk tRNA (Roche) from either *E. coli* (tRNA^Val^, tRNA^Lys^, tRNA^Gln^, tRNA^Cys^, tRNA^Met^) or yeast (tRNA^Arg^, tRNA^Trp^, tRNA^Glu^, tRNA^Lys^, tRNA^Thr^, tRNA^Leu^) via hybridization to immobilized complementary oligoDNAs presented in Table S1, as described previously^**30,38** – **40**^. More recently we have prepared bulk yeast-tRNA from fresh Red Star yeast cakes as described ^**41**^. ACE-tRNA^Arg^_UCA_ was prepared as described^**15**^. All tRNAs were charged with their cognate amino acids at 37°C using crude synthetase preparations from either *E. coli* or yeast which matched the source of the tRNA^**38,42,43**^. *S. cerevisiae* synthetase preparation was used to prepare Arg-ACE-tRNA^Arg^_UCA_. Atto 647-ε-lysine tRNA^Lys^, needed for the preparation of the Atto647-labeled Stop-POST5 complexes utilized in the fluorescence anisotropy termination assays, was prepared as described^**32**^. Cy5-heRF1 was prepared as described^**44**^ by inserting a short peptide tag ybbR (DSLEFIASKLA) between the TEV (Tobacco Etch Virus) protease cleavage site and the open reading Frame (ORF) using a QuickChange mutagenesis kit (Agilent) and then transformed into BL21(DE3) Codon Plus (Agilent) strain in the presence of ampicillin. Single colonies were grown overnight at 37 °C in 100 mL of LB-amp media. Overnight cultures were diluted to 0.1 A_600nm_ and grown to 0.6 A_600nm_ in 2L LB Amp media at 16 °C. 0.5 mM IPTG was added, and the cell cultures were incubated at 16 °C overnight. Cells were collected by centrifugation at 2700 x g (4000 rpm in a GS3 rotor) for 20 min at 4 °C. Cell pellet (∼5 g) was resuspended in eRF1/eRF3 Equilibration Buffer (100 mM HEPES-KOH, pH 7.4, 100 mM NaCl, 10% glycerol, 1 mM DTT, 50 mL) and lysed using a Qsonica sonicator at 30% for ten 15-second-pulse cycles with 30 s cooling on ice. Cell debris was spun down at 27 k x g (15 k rpm in an SS34 rotor) for 15 min at 4 °C. Cell lysate (∼ 50 mL) was loaded onto a 1.5 mL (3 mL slurry) TALON Superflow resin (Clontech) pre-equilibrated with equilibration buffer. The resin was washed three times with a 7.5 mL wash buffer (100 mM HEPES-KOH, pH 7.4, 10 mM imidazole, 100 mM NaCl, 10% glycerol, 1 mM DTT) and proteins were eluted in 0.5 to 1 mL fractions with elution buffer (100 mM HEPES-KOH, pH 7.4, 200 mM imidazole, 100 mM NaCl, 10% glycerol, 1 mM DTT). Fractions with ybbR tagged heRF1 were dialyzed against the storage buffer (100 mM HEPES-KOH, pH 7.6, 100 mM NaCl, 10% glycerol, 1 mM DTT) overnight and buffer exchanged into labeling buffer (50 mM HEPES, 10 mM MgCl_2_, and 1 mM DTT). For the labeling reaction, ybbR-heRF1 (10 µM, estimated using an e_280_ of 32110 M^-1^ cm^-1^) was incubated with CoA-Cy5 (SiChem, Bremen) at a 4:1 dye-to-protein ratio in the presence of 4 µM Sfp enzyme (a kind gift from the Christian Kaiser lab) at room temperature for 45 mins with continuous shaking. Excess dye was removed by passing the reaction mixture through a P-30 Bio Spin column (Bio-Rad). The labeled protein, having a labeling stoichiometry of ∼73%, was buffer exchanged into storage buffer (100 mM HEPES, 100 mM NaCl, 10% Glycerol, 1 mM DTT).

**Table 1.**
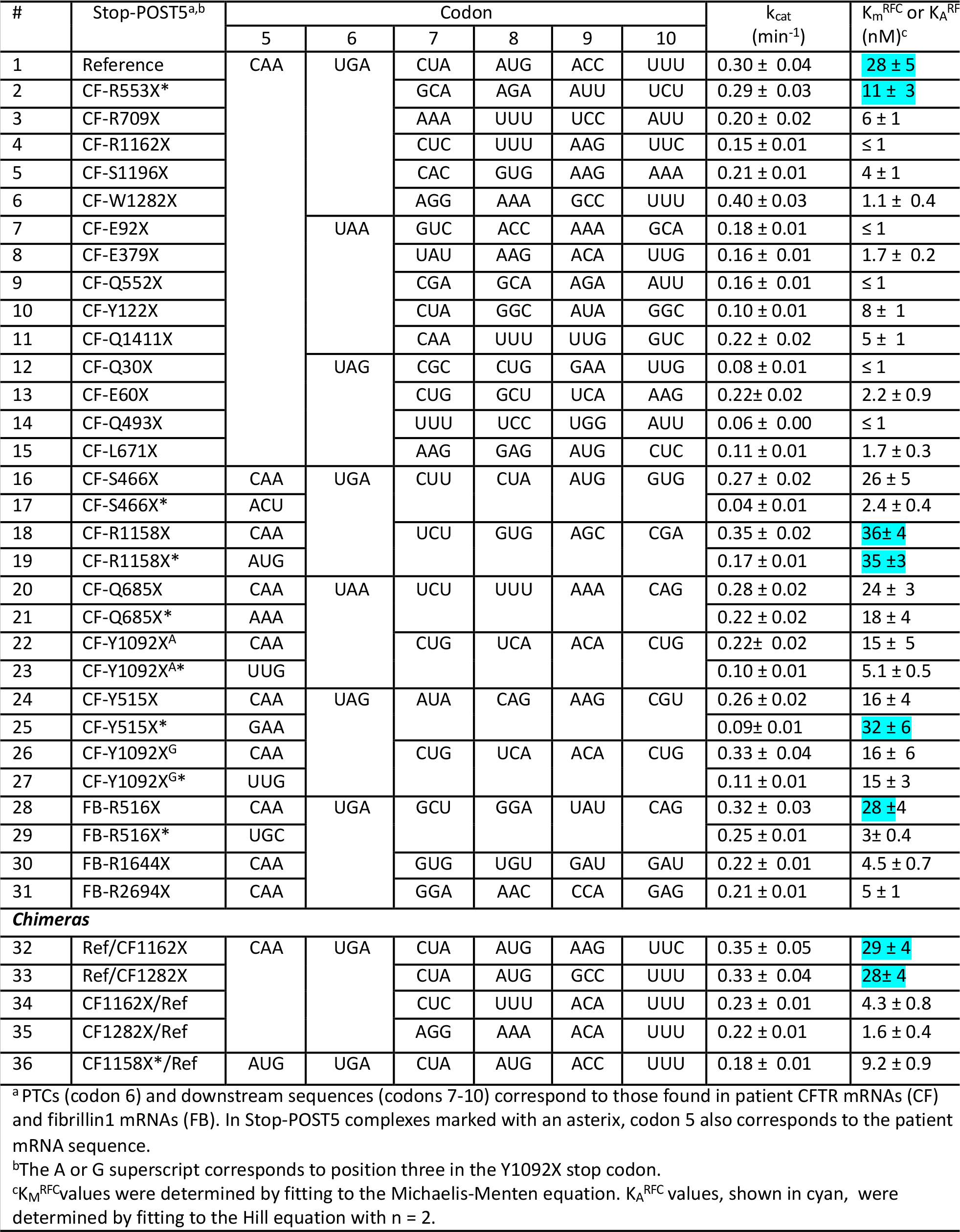
Termination Assay Results.

**Table 2.** Ataluren effects on Termination^a^ and Readthrough^b^.

| Table 2. Ataluren effects on Termination<sup>a</sup> and Readthrough<sup>b</sup> |  |  |  |  |  |  |  |  |
| --- | --- | --- | --- | --- | --- | --- | --- | --- |
| Stop-POST5 | Codons |  |  | Ata,<br>1 mM | Termination<br>Rate, (min<sup>-1</sup>) | K<sub>M</sub><sup>RFC</sup> or<br>K<sub>A</sub><sup>RFC,a</sup> | RE,<br>% | RE, %<br>+ 1 mM<br>Ata |
|  | 5 | 6 | 7 |  |  |  |  |  |
| CF-R1158X | CAA | UGA | UCU | -<br>+ | 0.15 ± 0.02<br>0.09 ± 0.03 | 36 ± 4 | 4 ± 1<br>14 ± 4 | 10 ± 4 |
| CF-R1158X* | AUG | UGA | UCU | -<br>+ | 0.08 ± 0.01<br>0.06 ± 0.01 | 35 ± 3 | 5 ± 3<br>12 ± 3 | 7 ± 4 |
| CF-Y515X* | GAA | UAG | AUA | -<br>+ | 0.07 ± 0.01<br>0.04 ± 0.01 | 32 ± 6 | 6 ± 3<br>32 ± 6 | 26 ± 6.7 |
| FB-R516X | CAA | UGA | GCU | -<br>+ | 0.20 ± 0.03<br>0.14 ± 0.01 | 28 ± 4 | 9 ± 5<br>20 ± 5 | 11 ± 7 |
| Reference | CAA | UGA | CUA | -<br>+ | 0.20 ± 0.02<br>0.08 ± 0.02 | 28 ± 5 | 2 ± 1<br>18 ± 2 | 16 ± 2.3 |
| CF-S466X | CAA | UGA | CUU | -<br>+ | 0.20 ± 0.01<br>0.11 ± 0.01 | 26 ± 5 | 6 ± 2<br>25 ± 6 | 19 ± 6 |
| CF-Q685X | CAA | UAA | UCU | -<br>+ | 0.13 ± 0.02<br>0.07 ± 0.02 | 24 ± 3 | 20 ± 2<br>60 ± 5 | 40 ± 5.5 |
| CF-Q685X* | AAA | UAA | UCU | -<br>+ | 0.18 ± 0.01<br>0.09 ± 0.02 | 18 ± 4 | 11 ± 2<br>30 ± 5 | 19 ± 5.4 |
| CF-Y515X | CAA | UAG | AUA | -<br>+ | 0.16 ± 0.02<br>0.07 ± 0.03 | 16 ± 4 | 4 ± 2<br>35 ± 4 | 31 ± 4.5 |
| CF-Y1092X<sup>G</sup> | CAA | UAG | CUG | -<br>+ | 0.18 ± 0.02<br>0.11 ± 0.01 | 16 ± 6 | 2 ± 1<br>27 ± 2 | 25 ± 2.2 |
| CF-G542X | CAA | UGA | GAA | -<br>+ | 0.18 ± 0.06<br>0.08 ± 0.04 | ND | 6 ± 3<br>24 ± 5 | 18 ± 5.8 |
| CF-Y1092X<sup>G*</sup> | UUG | UAG | CUG | -<br>+ | 0.12 ± 0.02<br>0.06 ± 0.01 | 15 ± 3 | 3 ± 1<br>26 ± 6 | 23 ± 6 |
| CF-Y1092X<sup>A</sup> | CAA | UAA | CUG | -<br>+ | 0.15 ± 0.02<br>0.08 ± 0.02 | 15 ± 5 | 5 ± 1<br>22 ± 3 | 17 ± 3 |
| CF-R553X* | CAA | UGA | GCA | -<br>+ | 0.18 ± 0.01<br>0.08 ± 0.01 | 11 ± 3 | ≤ 1 (0)<br>6 ± 4 | 6 ± 4 |
| CF1158X*/Ref | AUG | UGA | CUA | -<br>+ | 0.135 ± 0.003<br>0.095 ± 0.012 | 9.2 ± 0.9 | 3 ± 1<br>16 ± 2 | 13 ± 2 |
| CF-R709X | CAA | UGA | AAA | -<br>+ | 0.15 ± 0.02<br>0.07 ± 0.02 | 6 ± 1 | 4 ± 1<br>21 ± 1 | 17 ± 1.5 |
| CF-Y1092X<sup>A*</sup> | UUG | UAA | CUG | -<br>+ | 0.10 ± 0.01<br>0.06 ± 0.01 | 5.1 ± 0.5 | 12 ± 1<br>33 ± 6 | 21 ± 6 |
| FB-R2694X | CAA | UGA | GGA | -<br>+ | 0.16 ± 0.02<br>0.06 ± 0.02 | 5 ± 1 | 6 ± 3<br>20 ± 4 | 14 ± 5 |
| CF-Q1411X | CAA | UAA | CAA | -<br>+ | 0.16 ± 0.02<br>0.08 ± 0.02 | 5 ± 1 | 2 ± 1<br>13 ± 2 | 11 ± 2.2 |
| CF-S1196X | CAA | UGA | CAC | -<br>+ | 0.18 ± 0.01<br>0.15 ± 0.01 | 4 ± 1 | No RT (0)<br>2 ± 1 | 2 ± 1 |
| FB-R516X* | UGC | UGA | GCU | -<br>+ | 0.21 ± 0.03<br>0.14 ± 0.02 | 3.0 ± 0.4 | 0.9 ± 0.4<br>8.5 ± 2.3 | 7.6 ± 2.3 |
| CF-S466X* | ACU | UGA | CUU | -<br>+ | 0.05 ± 0.01<br>0.04 ± 0.01 | 2.4 ± 0.4 | 4 ± 2<br>6 ± 2 | 2 ± 2.8 |
| CF-L671X | CAA | UAG | AAG | -<br>+ | 0.10 ± 0.01<br>0.09 ± 0.01 | 1.7 ± 0.3 | 3 ± 2<br>3 ± 2 | 0 |
| CF-W1282X | CAA | UGA | AGG | -<br>+ | 0.39 ± 0.05<br>0.36 ± 0.03 | 1.1 ± 0.4 | 22 ± 6<br>25 ± 2 | 3 ± 6 |
| CF-R1162X | CAA | UGA | CUC | -<br>+ | 0.14 ± 0.01<br>0.14 ± 0.01 | ≤ 1 | < 1 (0)<br>1.2 ± 0.2 | 1.2 ± 0.2 |
| CF-E92X | CAA | UAA | GUC | -<br>+ | n.d<br>n.d | ≤ 1 | < 1<br>< 1 | 0 |
| CF-Q30X | CAA | UAG | CGC | -<br>+ | 0.09 ± 0.01<br>0.07 ± 0.02 | ≤ 1 | 2.7 ± 0.7<br>5.1 ± 1.1 | 2.4 ± 1.3 |

<sup>a</sup>K<sub>A</sub><sup>RFC</sup> values are shown in cyan

### Preparation of CrPV-IRES-variants

All CrPV-IRES-variant plasmid clones were obtained from Twist Biosciences and subsequently converted into the 309 residues long CrPV-IRES-mRNA by the HiScribe^®^ T7 high yield RNA synthesis kit (New England Biolabs, Catalog # E2040S) for use in the preparation of 80S.Stop-IRES and Stop-POST5 complexes. Shown below is the Reference CrPV-IRES-mRNA sequence. The colored residues mark the beginning of the attached mRNA moiety, with the green residues encoding the 5 amino acids upstream of the stop codon UGA, and the yellow residues encoding the 4 amino acids downstream from UGA.

GGAUCCUAAUACGACUCACUAUAGGGAGACCGGAAUUCAAAGCAAAAAUGUGATCUUGCUUGUAAAUACAAUUUUGAGAGGUUAAUAAAUUACAAGUAGUGCUAUUUUUGUAUUUAGGUUAGCUAUUUAGCUUUACGUUCCAGGAUGCCUAGUGGCAGCCCCACAAUAUCCAGGAAGCCCUCUCUGCGGUUUUUCAGAUUAGGUAGUCGAAAAACCUAAGAAAUUUACCUUUCAAAGUGAGACAAUGACUAAUGACAUUUCAAGAUACCAUGGAAGACGCCAAAAACAUAAAGAAAGGCCCGGAAGCUU.

Other CrPV-IRES-mRNA contexts, with sequences drawn mostly from CFTR (cystic fibrosis) and FBN1 (Marfan syndrome) patients harboring PTCs had the sequence design shown below.

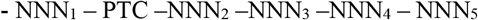

The NNN_1_ upstream codon (codon 5 in Table 1) was either held constant as CAA, encoding tRNA^Gln^_UUG_, or variable, matching what is found in the patient sequence. PTC (codon 6 in Table 1) is the stop codon (UGA/UAG/UAA) and NNN_2_ to NNN_5_ are downstream codons (codons 7 - 10 in Table 1) mostly reproducing patient sequences. All IRES sequences were verified by sequencing performed in the University of Pennsylvania Genomic and Sequencing Core Facility.

### Preparation of Stop-POST5 complexes

With one exception, all Stop-POST5 complexes were prepared by adding all 5 aminoacyl-tRNAs in a single step to CrPV-IRES-mRNA programmed 80S ribosomes in Buffer 4 (40 mM Tris HCl, pH 7.5, 80 mM NH_4_Cl, 5 mM Mg(Ac)_2_,100 mM KOAc, and 3mM β-mercaptoethanol) in the presence of eEF1A, eEF2, and GTP as previously described^**30,32,37**^. The exception is the Atto647-labeled CF-Q685X*Stop-POST5 complex used in termination studies, which contains a Lys residue at both codons 2 and 5 (#21, Table 1) and was prepared in two steps. First, by addition of Phe-tRNA^Phe^ and Atto647-ε-lysine tRNA^Lys^ in the presence of eEF1A and eEF2 and GTP, and isolation of the ribosome bound Atto647-labeled Phe-Lys-tRNA^Lys^ by sedimentation through a 1.1M sucrose cushion (120 min at 110,000 rpm using a S52-ST rotor on a Sorvall Discovery M120 SE ultracentrifuge). The resulting pellet was washed twice with Buffer 4 to remove any unbound Atto647-ε-lysine tRNA^Lys^, resuspended in Buffer 4, and centrifuged at 14,000 rpm for 30 min using a fixed angle F-45-30-11 rotor on an Eppendorf 5417C centrifuge. In the second step, Val-tRNA^Val^, Arg-tRNA^Arg^ and unlabeled Lys-tRNA^Lys^ were added in the presence of eEF1A, eEF2 and GTP to complete preparation of the Atto647-labeled CF-Q685X* Stop-POST5 complex.

### Termination assay

Termination assays were performed at 25 °C in Buffer 4 with a TECAN SPARK multimode plate reader equipped with a monochromator, either in a 96-well (96 well Greiner^®^) or a 384-well (Corning) black flat bottom plate as described ^**32, 37**^. Two solutions were prepared in Buffer 4, one containing 0.05 μM Atto(pep)-Stop-POST5 and 1 mM GTP and the other containing eRF1 (0-0.12 μM), eRF3 (0.8 μM) and 1 mM GTP in Buffer 4. Reaction was initiated by mixing the two solutions. All concentrations listed are final after mixing. The decrease in fluorescence anisotropy on release of the Atto647-labeled pentapeptide from the ribosome following eRF1-catalyzed hydrolysis of ribosome-bound pentapeptidyl-tRNA was monitored continuously. Excitation and emission wavelengths are preset in the plate reader for Atto-647. For measurements involving ataluren, both the RFC and Atto(pep)-Stop-POST5 solutions contained 1 mM ataluren. Time dependent decreases in fluorescence anisotropy were analyzed with Graphpad Prism, using the single exponential with an asymptote model to obtain peptide dissociation rates for all Stop-POST5 complexes except for CF-W1282X, the results for which required a biphasic equation. The fast phase, corresponding to about 50% of the release, was used to quantify the peptide release rate. All rate data were fit to the Michaelis-Menten or Hill equations to determine values of k_cat,_ and K_MRFC_ or KA ^RFC^.

For most of the mRNA sequences, termination rate, k_obs_, plotted *vs*. free RFC concentration (RFC_free_), was well fit by the Michaelis-Menten equation. yielding maximum dissociation rate, k_cat_ and the half-saturation concentration of free RFC, KM^RFC^. For 8 of the sequences, marked in cyan in Table 1, the plots of k_obs_ *vs*. RFC_free_ were S-shaped and were fit by the Hill equation (Eq. 1), with *n* = 2, yielding k_cat_ and K_A_^RFC^.

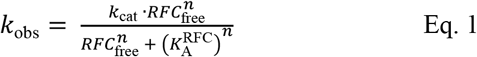

The Hill equation reduces to the Michaelis-Menten equation when *n* = 1 (Eq. 2)

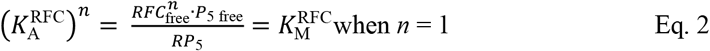

*RFC*_free_ was calculated from the known total concentrations of RFC (i.e., *RFC*_total_) and of Stop-POST5 complex, shown as *P*_5 total_ in Eq. 3. *RP*_5_ is the concentration of the RFC·Stop-POST5 complex.

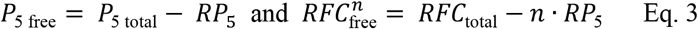

For *n* = 1, combining these equations into one relationship yields a quadratic equation which was solved analytically for *RFC*_free_. For *n* = 2, combining the equations yields a cubic equation which was solved numerically for *RFC*_free_ using the “Solver” plug-in in Excel. *k*_cat_ and *K*M^RFC^ or *K*A^RFC^ were adjusted to fit Eq. 1 to the termination rate data *k*_obs_ *vs. RFC*_free_ with *n* = 1 or 2. All measurements were repeated three times. The data shown are averages of 3 measurements ± SD.

Readthrough assays (RE assays)

In RE Assay 1, described earlier^**31**^, we use co-sedimentation to determine the stoichiometry of [^3^H]-Stop-Pre6 complex formed when Stop-POST5 complex (0.05 µM) is incubated for 2 minutes @ 25 °C, in Buffer 4, sufficient for essentially complete reaction, with [^3^H]-labeled suppressor aa-tRNA (0.2 µM), either near-cognate or cognate to the stop triplet in codon 6, in the presence of eEF1A (1 µM) and GTP (1 mM) and the absence or presence of eRF1 (0.05 µM) and eRF3 (0.2 µM). Near cognate suppressor tRNAs, Trp-tRNA^Trp^_CCA_,Tyr-tRNA^Tyr^_GUA_ and Gln-tRNA^Gln^_UUG_, added at 0.2 µM concentration, were used to measure readthrough at stop codons, UGA, UAG and UAA, respectively. Reaction was initiated by mixing solution A containing Stop-POST5 complex with solution B containing TC ± RFC. All concentrations listed are final after mixing. When TRIDs were added, both solutions contained ataluren and G418, added either singly or in combination, at their final concentrations. Reactions were quenched with ice-cold 100 μL of 0.5 M MES buffer (pH 6.0) and placed on ice, 100 pmol of 70S carrier ribosomes were added, and ribosomes were collected by ultracentrifugation as previously described^**31**^. In RE Assay 2, performed using a plate reader to measure fluorescence quantum yield increase during conversion of a proflavin-labeled Stop-POST5 complex to a proflavin-labeled Stop-POST6 complex. In this assay, the [^3^H]-labeled suppressor aa-tRNA is replaced by a proflavin-labeled suppressor aa-tRNA (0.1 µM), prepared as described^**31**^, with either tRNA^Trp^_CCA_ or tRNA^Gln^_UUG_, having labeling stoichiometries of 2 and 1, respectively. All other conditions of reaction are as in RE Assay. 1. After mixing solution A with solution B, fluorescence was measured on the TECAN SPARK multimode reader for 10 minutes by exciting the fluorophore at 462 nm, and monitoring emission at 515 ± 15 nm. The value of ΔF was taken at 2 min, which corresponded to full reaction. The two assays gave similar results (Figure 2C,D).

**Figure 2.**
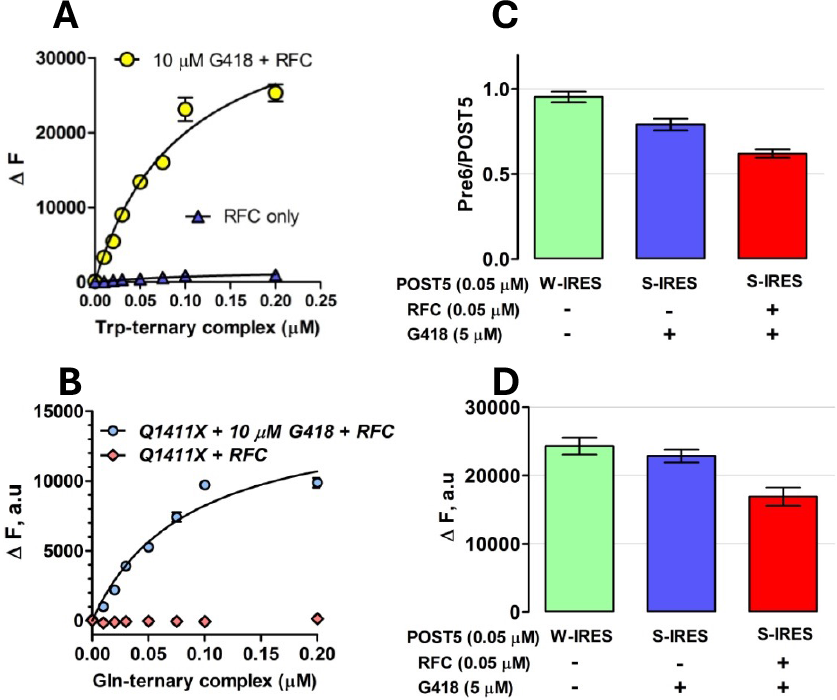
Readthrough Assays. **A,B.** Titration of proflavin-labeled suppressor aa-tRNA, with either tRNA^Trp(CCA)^ for UGA stop codon in Reference Stop-POST5 (Table 1, #1) and tRNA^Gln(UUG)^ for UAA stop codon in CF-Q1411X (Table 1, #11). **C,D**. Comparing readthrough assay results. S-IRES refers to the Reference Stop-POST5 complex. W-IRES refers to a similar complex but with UGG, cognate to tRNA^Trp^ replacing UGA at codon 6. **C**. Assay 1, measuring [^3^H] co-sedimenting with POST6 complexes. **D**. Assay 2, measuring fluorescence increase, ΔF, accompanying formation of POST 6 complexes. Note that, in the absence of RFC, G418 induces almost complete readthrough of S-IRES, as previously demonstrated.^**31**^

## RESULTS

### Effects of sequence context on termination

#### Ensemble experiments

In our earlier use of PURE-LITE ^**31,32,37,38**^, we prepared a pretermination complex, denoted here as Reference Stop-POST5, having a UGA stop codon at position 6, an upstream CAA codon at position 5, and the downstream sequence CUA AUG ACC UUU at codons 7-10 encoding L, M, T, F (Fig. 1A). The UGA stop codon, the upstream CAA codon, and the downstream CUA codon were chosen to favor readthrough^**38**^. The relatively high (20%) basal readthrough by near-cognate Trp-tRNA^Trp^ permitted us to examine both termination and readthrough reactions. Readthrough was measured at codon 6 or 8 by incorporation into the nascent peptide of either [^3^H]-Trp or [^35^S]-Met, respectively. Here we extend these studies to first examine the effects on termination of varying **i**. stop codon identity, **ii**. four codons downstream of the stop codon, and **iii**. one codon upstream of the stop codon, which, based on studies referred to above^**23-25**^, include the mRNA contexts most consequential in determining REs. These variations necessitated the construction of a significant number of Stop-POST5 complexes, which, compared to Reference-Stop-POST5, can have changes in codons 5 – 10, as indicated in Table 1. The large majority of the Stop-POST complexes examined, 29/36 in Table 1, maintained CAA as the immediate codon upstream of the stop codon, to highlight differences due to downstream sequences, reflecting their known greater influence on RE^**23-25, 28**^.

We used a plate-reader assay^**32,37**^ to determine the rates of termination for each of the Stop-POST5 complexes as a function of RFC concentration. This assay measures fluorescence anisotropy decrease as a fluorescent-labeled nascent peptide is released from the Stop-POST5 complex by the catalytic activity of RFC. The termination results for all Stop-POST5 complexes are presented in Table 1. The results for 28 of these Stop-POST5 complexes show a rectangular hyperbolic dependence on RFC concentration, corresponding to an apparent single effective site of RFC binding to the ribosome, fitting well to the Michaelis-Menten equation, and allowing quantification of KM^RFC^. In contrast, 8 Stop-POST5 complexes display a clear S-shaped dependence on RFC concentration, providing evidence for positive homotropic cooperativity in binding of RFC to the ribosome. KA^RFC^ values were determined for these sequences from the Hill equation (eq. 1-see Materials and Methods), after calculating free [RFC], taking into account RFC bound, and setting Hill *n* = 2.

Our lower limit for estimation of either KM^RFC^ or KA^RFC^, ≤1.0 nM, is determined by the lowest concentration of Stop-POST5 complex we employed which gave a reliable fluorescence anisotropy decrease. Within this limitation, KM^RFC^ and KA^RFC^ values were broadly distributed over a ≥35 fold range, with all three of the stop codons. In contrast, k_cat_ values vary over a narrower range, about 10-fold, and are not broadly distributed, with 90% falling between 0.1 - 0.3 min^-1^. Termination rates vs. [RFC]_free_ for representative sequences are presented in Figure 3. Four sequences for which the upstream codon (position 5, nts -1 to -3 relative to the stop codon) CAA encoding Gln is held constant (#s 9, 16, 20 and 22 in Table 1), show the clear dependence of KM^RFC^ on the downstream codons 7 – 10: ≤ 1 nM for # 9 (Figure 3A) and higher for #s 22 (15 ± 5 nM), 20 (24 ± 3 nM), and 16 (26 ± 5 nM) (Figures 3B, 3C, and 3D, respectively).

**Figure 3.**
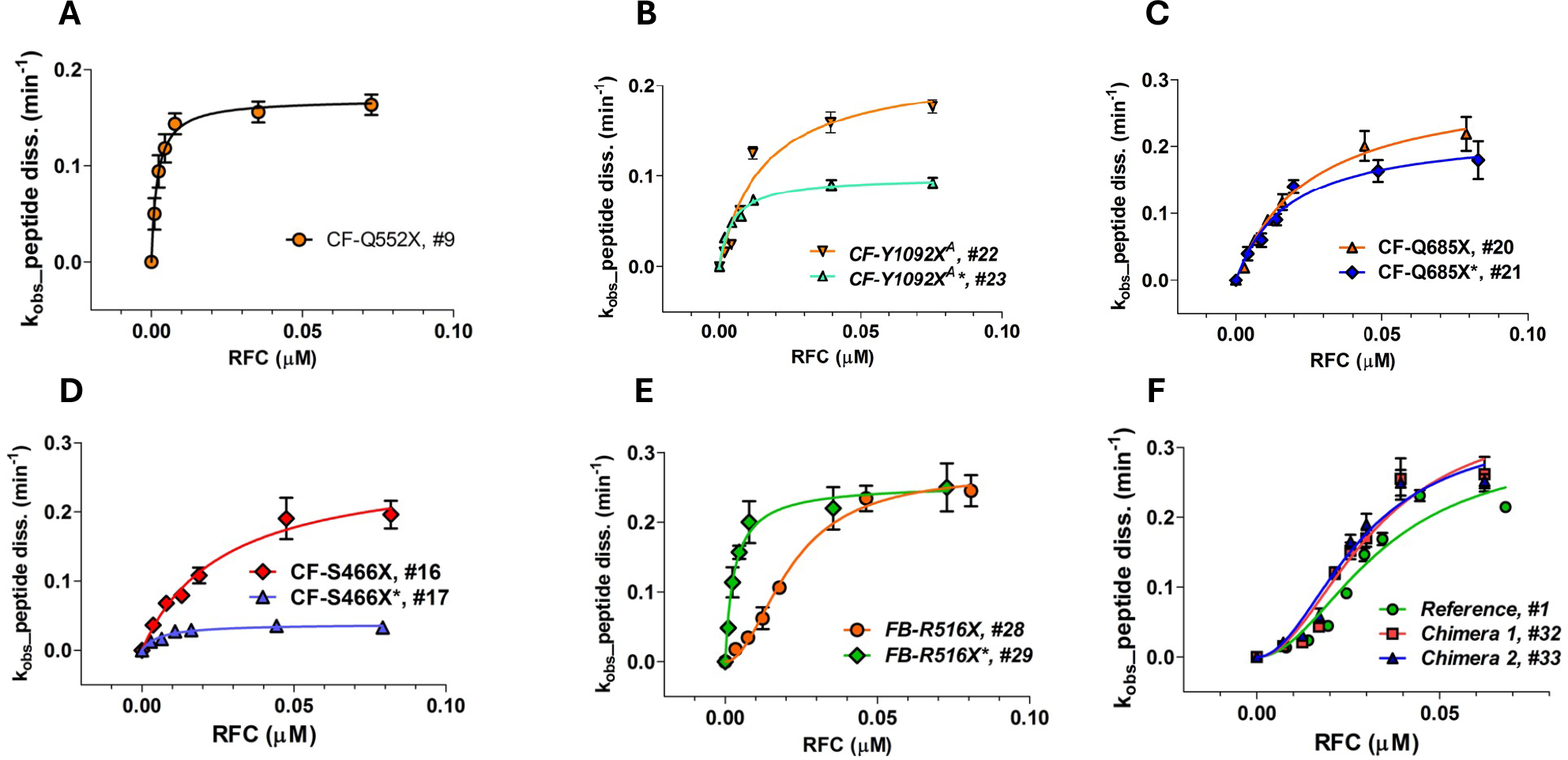
Representative results showing the dependence of the rate of peptide dissociation from Stop-POST5 complexes, numbered as in Table 1, on the free concentration of RFC. **A – D.** Results fit to the Michaelis-Menten equation. **B – D**. Variable effects due to changing the upstream codon 5 from CAA to the codon present in a patient mutation are shown. **E**. Results fit to the Hill equation (#28) or the Michaelis-Menten equation (#29). **F**. Results fit to the Hill equation, n=2, for Reference (#1, green), Chimera 1 (#32, red), and Chimera 2 (#33, blue).

Changing the upstream codon from CAA, which generally favors readthrough^**23**^, to the codon present in each patient sequence, has variable effects (Table 1). For the seven pairs of Stop-POST complexes examined, k_cat_ either shows little change (sequence #s 20, 21; 28,29) or substantial decrease for the disease sequences (2-to 6-fold, #s 16, 17; 18, 19; 22, 23; 24, 25; 26, 27). Greater variability is seen in the interaction of RFC with Stop-POST5 as measured by either KM^RFC^ or K_A_^RFC^, which can show little change (#s 18,19; 20,21; 26,27), significant decrease (3-to 10-fold: #s 16, 17; 22, 23; 28, 29), or significant increase (2-fold, #s 24, 25). Interestingly, replacing the upstream CAA at codon 5 with the patient codon can result in a change in RFC concentration dependence from S-shaped to a rectangular hyperbola (Figure 3E, #29 vs. #28) or vice-versa (#25 *vs*. #24). In order to determine the relative effects on termination activity of downstream codons 7 and 8 *vs*. 9 and 10 we examined Stop-POST5 complexes derived from four chimeric mRNAs. In Ref/CF-1162X (#32) and Ref/CF1282 (#33), codons 7 and 8 are derived from Reference Stop-POST5 (#1), which has an S-shaped dependence on RFC concentration and a K_A_^RFC^ of 28 ± 5 nM. In contrast, codons 9 and 10, derived from CF-1162X (#4) and CF-1282X (#6), respectively, catalyze peptide release with a rectangular hyperbolic RFC concentration dependence, with K_M_^RFC^ values of ≤ 1 nM. In CF1162/Ref (#34) and CF1282/Ref (#35), the identities of the downstream codons are reversed. The termination activity results demonstrate clearly that the identities of the proximal downstream codons 7 and 8 largely determine the response of Stop-POST5 complexes to RFC, whereas the more distal downstream codons 9 and 10 matter less. Thus, as shown in Figure 3F and Table 1, the dependence of termination on RFC concentration of the Reference (#1), Ref/CF-1162X (#32) and Ref/CF1282 (#33) Stop-POST5 complexes are almost identical with respect to both K_A_^RFC^ and k_cat_ whereas K_M_^RFC^ values for CF1162/Ref (#34) and CF1282/Ref (#35) are much closer to those found for CF-1162X (#4) and CF-1282X (#6), respectively, than to that found for Reference K_A_^RFC^.

#### Single molecule Fluorescence Resonance Energy Transfer (smFRET) experiments

RFC binding to a pretermination complex such as Stop-POST5 is followed by rapid release of eRF3 and by a much slower release of eRF1^**45**^. We employed an smFRET approach to determine whether the large range of K_m_^RFC^ and K_A_^RFC^ values as a function of sequence context shown in Table 1 is due either to differences in the rate of RFC binding to Stop-POST5 complexes (k_arrival,app_) or, following peptidyl-tRNA hydrolysis, in the rate of release from the ribosome of tRNA (k_dis,tRNA,app_) or eRF1 (k_dis,heRF1,app_). For this purpose we prepared a Cy5-labeled derivative of human eRF1 (Cy5-heRF1) and used it to form an active RFC complex with heRF3.GTP (Cy5-heRFC). Binding of Cy5-heRF1 to Stop-POST5 complexes, each of which contained tRNA-labeled FKVRQ-tRNA^Gln^(Cy3) bound in the P-site, adjacent to the stop codon, generated a transient FRET signal which disappears following peptidyl-tRNA hydrolysis and release of Cy5-eRF1 and tRNA^Gln^(Cy3). In these experiments, the Cy3-labeled Stop-POST5 complex was attached to a microscope cover slip chamber via the mRNA and a 3’ a biotin-streptavidin linkage (Figure 4A). To initiate the binding reaction, Cy5-heRFC (35 nM) was injected into the flow cell 10 seconds after the video recording began (Figure 4B). The arrival of Cy5-heRFC, detected directly using alternating laser excitation (ALEX) at wavelengths of 532 nm and 640 nm (panel ii, red trace) was closely followed by partial quenching of the Cy3-tRNA fluorescence (panel i, green trace) and an increase in the sensitized emission of Cy5 (panel i, red trace) due to FRET. The transient FRET efficiency of *E* = ∼0.25 between Cy5-heRF1 and Cy3-tRNA (panel iii, blue trace) is consistent with heRF1 being successfully accommodated within the ribosomal A site, allowing peptidyl-tRNA hydrolysis to proceed. Cy5-eRF1 and tRNA^Gln^(Cy3) were released with some delay after hydrolysis.

**Figure 4.**
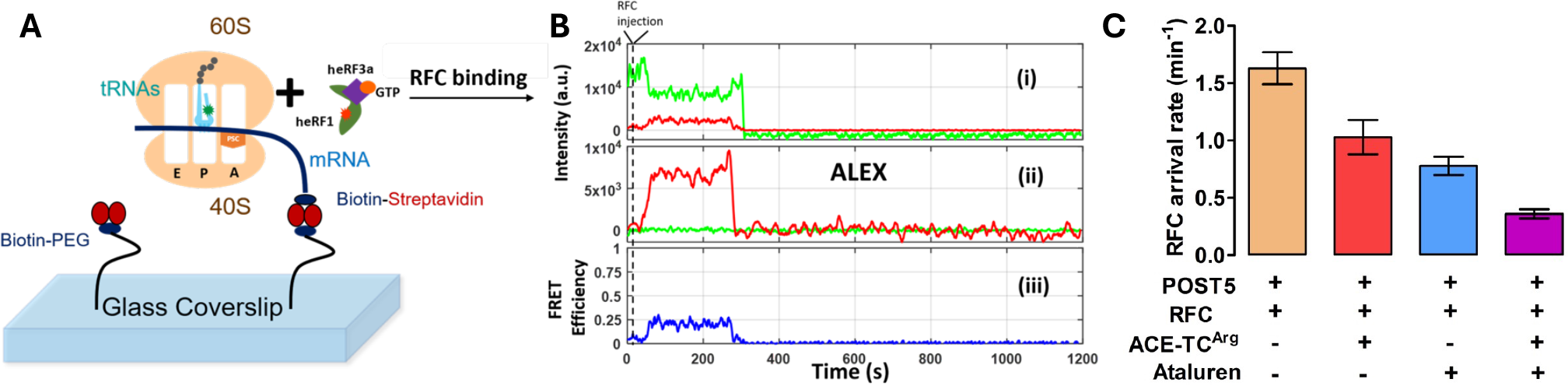
Sm FRET measurement of RFC interaction with Stop-POST 5 complexes. *A*. Experimental design. ***B***. Sample traces measuring the time dependence for interaction of an RFC containing Cy5-heRF1 with Reference-Stop-POST5 complex containing FKVRQ-tRNA_Gln_(Cy3) bound in the P-site. These traces permit measurement of the rates of RFC arrival (k_arrival,app_), dissociation of tRNA_Gln_(Cy3), and dissociation of Cy5-heRF1 (k_dis,tRNA,app_) from, respectively, the times of ALEX (ii) or FRET (iii) appearance, Cy3 disappearance (i) and ALEX disappearance (ii). These values for 13 Stop-POST5 complexes are presented in Table S2. ***C***. Effects of ataluren and ACE-tRNA, added separately or in combination, on k_arrival,app_.

We performed the smFRET experiment on 13 different Stop-POST5 complexes. Apparent rate constants for RFC binding (k_arrival,app_) and both tRNA^Gln^ (k_,tRNAGln,app_) and heRF1 (k_dis,heRF1,app_) release for each of the Stop-POST5complexes, measured at a single RFC concentration (35 nM) are shown in Table S2. All k_arrival_,_app_ values fall within a 2-fold range, as do 12 of the 13 values of k_,tRNA,app_ and k_dis,heRF1,app_, contrasting sharply with the ≥ 30-40 fold difference in measured K_M_^RFC^ and K_A_^RFC^ values for the same Stop-POST5 complexes. One exception to the limited range of these data is CF-R553X*, for which k_dis,tRNA,app_ and k_dis,heRF1,app_ are 4-to 5-fold slower than the fastest ones. These results provide a clear indication that differences in K_M_^RFC^ and K_A_^RFC^ are not due to differences in the rates of RFC binding or of tRNA or eRF1 release. Possible later reaction steps which control termination rates and RFC activity are reversible RFC dissociation, eRF3 GTPase, P_i_ release, or eEF3.GDP dissociation. Further studies will be needed to determine which of these steps show a clear correlation with K_M_^RFC^/K_A_^RFC^ values.

### Effects of sequence context on readthrough efficiency (RE), as modified by addition of ataluren

RE measured in the absence of nonsense suppressors reflects a competition between RFC and near-cognate TC for covalent reaction with Stop-POST5, i.e., hydrolysis of pentapeptidyl-tRNA *vs*. elongation to hexapeptidyl-tRNA (Figure 1B). We use two different assays to determine RE, which give equivalent results (Figure 2C,D). Ataluren addition, by competitively inhibiting RFC binding^**32**^ to Stop-POST5 without affecting near-cognate TC binding^**31**^, alters the RFC *vs*. TC competition in favor of the near-cognate TC, thereby stimulating readthrough. Ataluren effects, added at 1 mM, on termination and readthrough for 27 Stop-POST5 complexes are presented in Table 2. ΔRE values (± ataluren) generally increase as K_M_^RFC^/K_A_^RFC^ values increase, indicating weaker interaction of RFC with a Stop-POST5 complex, consistent with our expectations based on ataluren competitive inhibition of RFC. The Spearman correlation coefficient between these parameters is r = 0.64 (Figure 5A). Because ΔRE values are consistently lower for K_A_^RFC^ values than for K_M_^RFC^ values, we observe much stronger correlation (Spearman r = 0.89) between readthrough and termination effects when the 21 K_M_^RFC^ values are considered separately (Figure 5B).

**Figure 5.**
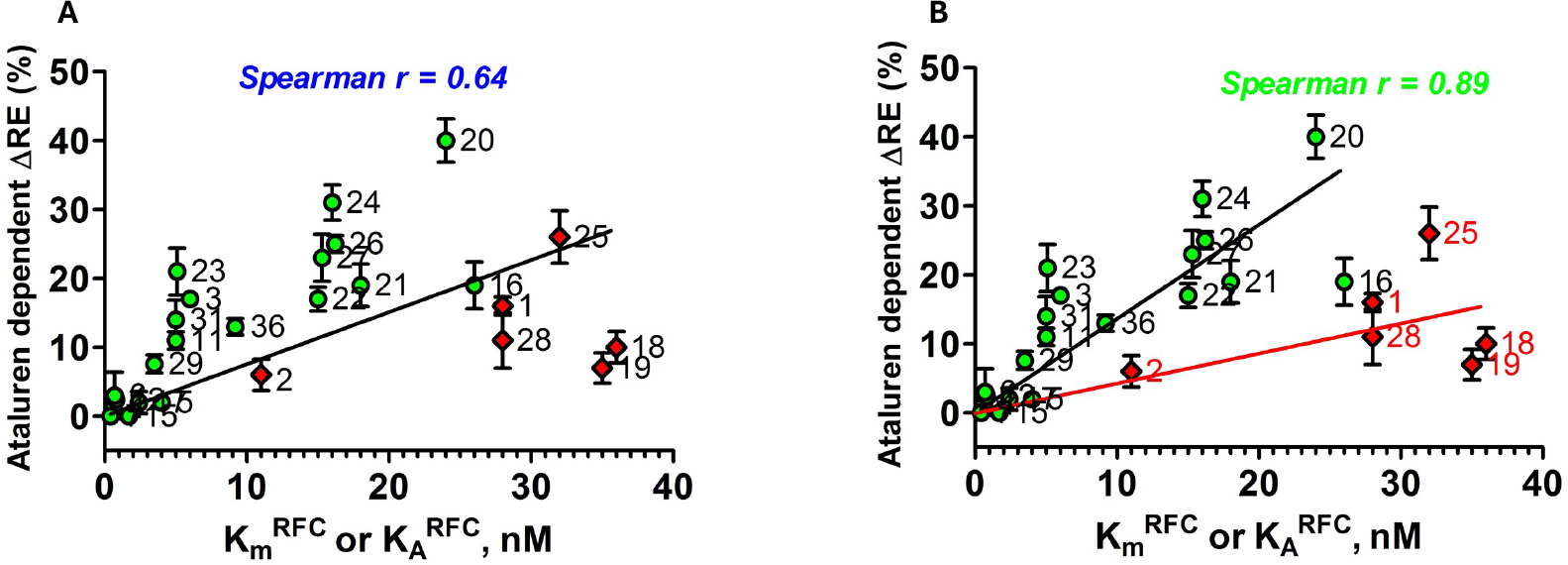
Correlation of ataluren stimulation of RE and K_M_^RFC^ (green-filled circles) and K_A_^RFC^ (red-filled diamonds) values. Stop-POST5 complexes are numbered as in Table 1. **A.** All values are fit to a linear regression model. **B**. K_M_^RFC^ and K_A_^RFC^ values are fit separately to linear regression models.

The ΔRE results in Table 2 were obtained at single concentrations of both ataluren (1 mM) and Suppressor-TC (Sup-TC, 0.2 µM), raising the possibility that outliers might arise from Stop-POST5 complexes differing in their dependencies on ataluren and Sup-TC concentrations. This does not appear to be the case, based on results presented in Figures S1A and S1B, showing that Stop-POST5 complexes having similar K_M_^RFC^/K_A_^RFC^ values but differing in ΔRE values have similar dependencies on ataluren and Sup-TC concentrations.

### Effects on readthrough efficiency of adding ataluren in combination with G418

Added separately, both ataluren and G418 stimulate readthrough of Reference Stop-POST5 complex^**31**^, leading to formation of FKVRQW-tRNA^Trp^. Combining ataluren and G418 leads to increased readthrough, as expected based on their orthogonal mechanisms for stimulating RE^**31**^. Sample results presented in Figure 6 show that effects of these combinations of nonsense suppressors can be additive (Stop-POST5 complexes Reference, CF-G542X, CF-R709X, CF-S446X) or synergistic (Stop-POST5 CF-R553X*). Neither CF-E92X nor CF-R1162X show ataluren stimulation of G418-induced readthrough, consistent with their tight RFC binding and resistance of these Stop-POST5 complexes to readthrough stimulation by ataluren alone.

**Figure 6.**
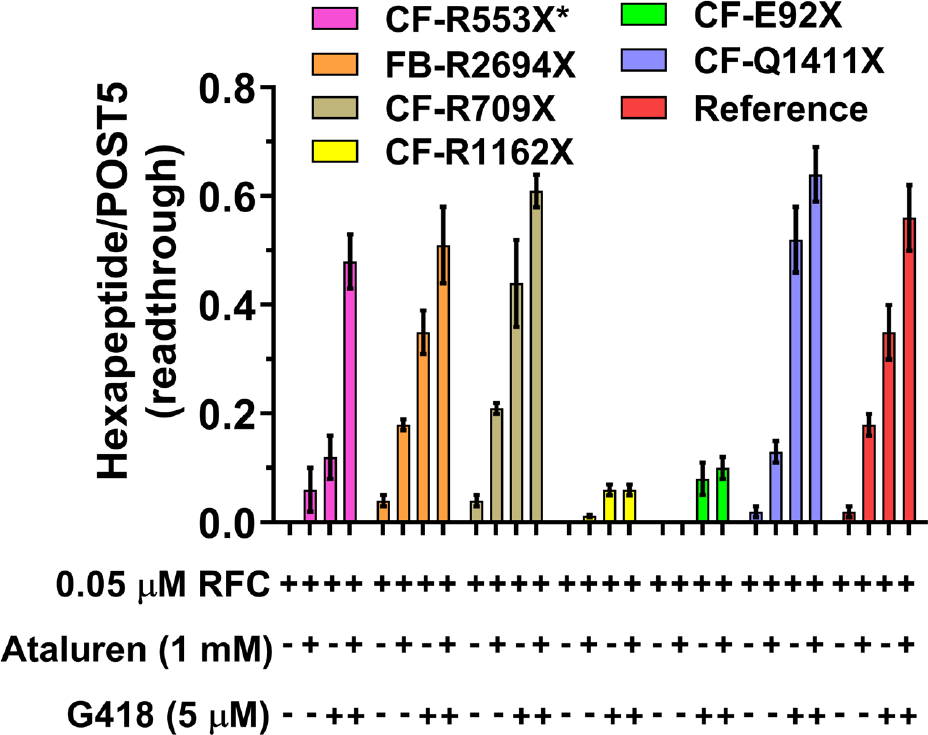
Ataluren and G418 promote readthrough of CFTR and FBN1 mutants either synergistically (CF-R553X*) or additively (4 others). Ataluren has little or no effect in promoting readthrough of CF-R1162X or CF-E92X

### Effects on termination and readthrough efficiency of adding ataluren in combination with ACE-tRNA^**Arg**^_**UCA**_

Since TC is a molecular mimic of RFC^29^, competing directly with it for binding to the A-site of the pretermination Stop-POST5 complexes used in this work (Figure 1B), we expect that ACE-tRNA^Arg^_UCA_ will compete with RFC for binding to Stop-POST5 complexes with a UGA stop codon, leading to both inhibition of termination and directly forming the readthrough product. It has been proposed^**46**^ that the identity of the peptidyl-tRNA bound in the P-site of a pretermination complex could affect this competition. To test this point, we determined ensemble termination rates and REs for seven different Stop-POST5 complexes in the presence of ACE-tRNA^Arg^_UCA_ and ataluren, added separately or in combination, giving the results displayed in Figures 7A and 7B. In these experiments the ACE-tRNA^Arg^_UCA_ TC and RFC concentrations were chosen to make clear the effects of ataluren on the competition between these two complexes. All seven Stop-POST5 complexes, containing P-site bound FKVRZ-tRNA^Z^ (Z = Q, T, or M) gave consistent results between the termination and readthrough assays. Six sequences showed additive effects between ataluren and ACE-tRNA^Arg^_UCA_ both in inhibiting RFC activity and stimulating readthrough in the presence of RFC, leading to the formation of ribosome-bound FKVRZR-ACE-tRNA^Arg^_UCA_ (Z= Q or T). One sequence, CF-R1158X* (Z = M, Table 1, #19), resisted both ataluren (Table 2) and ACE-tRNA^Arg^_UCA_ inhibition of RFC activity (Figure 7A), and ACE-tRNA^Arg^_UCA_ readthrough of the UGA stop codon (Figure 7B). CF-R1158X* Stop-POST5 differs from Reference Stop-POST5 (Table 1, #1) in both its upstream codon 5 (AUG vs. CUA) and its downstream codons, 7 -10 (UCU GUG AGC CGA vs. CUA AUG ACC UUU). The lack of responsiveness to ACE-tRNA^Arg^_UCA_ appears to be due mainly to the difference in the downstream sequence, since the chimeric CF-R1158X*/Ref Stop-POST5 (Table 1, #36), having the variable sequence AUG UGA CUA AUG ACC UUU, displays considerable ACE-tRNA^Arg^_UCA_ and ataluren inhibition of RFC activity (Figure 7A) as well as ACE-tRNA^Arg^_UCA_ readthrough of the UGA codon and ataluren stimulation of such readthrough (Figure 7B).

**Figure 7.**
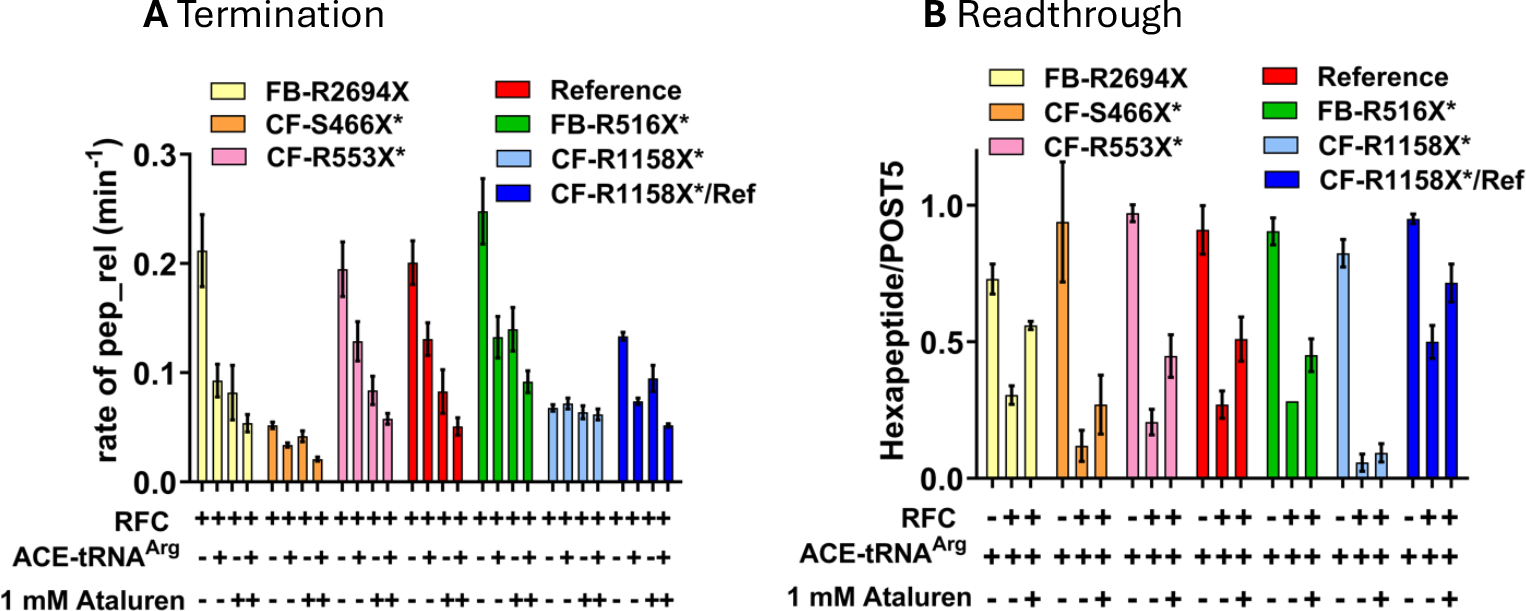
Ataluren acts consistently in its effect on ACE-tRNA^Arg^_UCA_ competition with RFC, as measured by termination **A** or readthrough **B**. The six Stop-POST5 complexes showing additive effects of ataluren and ACE-tRNA^Arg^_UCA_ in inhibiting termination **A**, also show partial reversal by ataluren of RFC inhibition of readthrough by ACE-tRNA^Arg^_UCA_ **B.** Similarly, neither ACE-tRNA^Arg^_UCA_ nor ataluren show significant effects in either inhibiting termination or stimulating readthrough for the CF-R1158X* Stop-POST5 complex. Concentrations, ACE-tRNA^Arg^_UCA_ (0.2 µM), RFC (0.0625 µM).

We applied the smFRET approach described above (Figure 4A,B) to determine the effects of adding ataluren and ACE-tRNA^Arg^_UCA_ on the interaction of Reference-Stop-POST5 with RFC. k_arrival,app_ is substantially decreased by each of these agents added alone, and is decreased significantly further when the agents are combined (Figure 4C), consistent with the ensemble results presented in Fig 7A. Neither agent has significant effects on k_heRF1 dis_, but each decrease k_tRNAGlnUUG_ by approximately 40%. However, in this case the effects are not additive.

## DISCUSSION

Here we demonstrate that PTC identity and mRNA sequence context modulate the catalytic activity of RFC in terminating peptide elongation, and, in so doing, determine the effectiveness of ataluren, a TRID which acts exclusively as a competitive inhibitor of RFC binding^**31,32**^, in stimulating readthrough efficiency (RE). We see such stimulation by ataluren acting alone, or cooperatively in combination with either the aminoglycoside G418 or an ACE-tRNA. RE reflects the competition between productive binding of a suppressor TC (Sup-TC) and RFC to an A-site stop codon. This competition strongly favors RFC in general, since natural Sup-TCs, which are only near-cognate to the stop codons, bind more weakly than RFCs and, once bound, are subject to rejection by proofreading, which reduces the frequency of productive binding. Ataluren shifts the Sup-TC *vs*. RFC competition in favor of the Sup-TC (Table 2), resulting in a significant correlation between sequence context effects on K_M_^RFC^/K_A_^RFC^ and ataluren stimulation of RE (Figure 5A), which is considerably more marked when K_M_^RFC^ results are considered separately (Figure 5B).

In contrast to these very clear relationships, our results raise three interesting questions which will require further experiments to resolve. First, why do most of the Stop-POST5 complexes we studied (Table 1) show a hyperbolic dependence of termination rate on RFC concentration, consistent with RFC binding to the single, structurally well characterized site on the ribosome leading to termination^**29,47**^, while a minority show a sigmoidal dependence, indicating that RFC binding to an additional site (or sites) is required for their termination activity? The K_A_^RFC^ (sigmoidal curves) and K_M_^RFC^ (hyperbolic curves) have mean values of 28 nM and 9 nM, respectively (Table 2), suggesting that weaker interaction of RFC with the Stop-POST5 complex could be a necessary, although not sufficient condition for sigmoidal dependence. An hypothesis which would account for our results is that some RFC complexes which interact weakly with the well characterized RFC site on the ribosome have suboptimal orientations for termination activity which RFC binding to an additional ribosomal site (or sites) can correct, and binding to such sites is not influenced by ataluren. The P1,P2 stalk of the ribosome could provide a possible location for such a site (or sites), to which one or more RFCs could bind via the eRF3 GTPase component of RFC^**48**^.

Second, what accounts for the clear deviations from strict linearity in the plot of ΔRE vs. K_M_^RFC^ (Figure 5B), most evident in the 2-fold ranges in ΔRE values for the Stop-POST5 complexes clustered at K_M_^RFC^ values between 5 -7 nM and 15-19 nM? We think it likely that these deviations arise from steps in the overall process of termination following Sup-TC binding to the ribosomal A-site which may not be sensitive toward ataluren. For example, it is clear from results presented in Figure 3 that the identity of the P-site bound peptidyl-tRNA can strongly influence ΔRE, consistent with results of others^**23-25,49**^. Also of possible relevance are results of Kolakada et al.^**50**^ showing that the identity of P-site bound peptidyl-tRNA affects nonsense-mediated decay (NMD), which also competes with readthrough in cells. In addition, using an smFRET approach we have recently shown that addition of the near-cognate Trp-TC to either the Reference Stop-POST5 complex (Table 1), or to a POST5 complex in which the UGA codon of Reference Stop-POST5 complex is replaced by the Cys codon UGU, requires several brief near cognate Trp-TC binding (sampling) events prior to stable binding leading to FKVRQW-tRNA^Trp^ formation^**51**^. Further, during the sampling period, a metastable ribosome conformation is formed, in which the accommodation step resulting in peptide elongation is retarded. Accumulation of this conformation may lead to ribosome stalling and the limited stoichiometry of peptide elongation observed when ribosomes interact with near-cognate TCs or inhibitory codon pairs^**31,52**^, and could certainly impact ΔRE values.

Third, how general a phenomenon is the resistance demonstrated by CF-R1158X*-Stop-POST5 to readthrough by ACE-tRNA^Arg^_UCA_ (Figure 7B)? As discussed above, the stop codon and the immediate upstream codon and 1-2 downstream codons are considered to be the most consequential for nonsense suppressor stimulated readthrough^**23-25**^. In CF-R1158X*-Stop-POST5 this quartet of codons 5 - 8 has the sequence AUG UGA UCU GUG (Table 1, #19). The readthrough results obtained with CF-R1158X*/Ref-Stop-POST5 (Figure 7B), which has the corresponding sequence AUG UGA CUA AUG (Table 1, #36), suggests that it might be the downstream UCU GUG codons which somehow reduce ACE-tRNA^Arg^_UCA_ readthrough efficiency, at least in this sequence, but perhaps more generally.

## CONCLUSION

Ataluren has been investigated for its potential to readthrough disease causing PTCs in clinical trials, animals, and cell-based assays, with variable outcomes^**9**^. The results presented in this paper (Tables 1 and 2 and Figures 3, 5-7), demonstrate the strong dependence on mRNA sequence context of ataluren-induced readthrough of PTCs, which is in turn strongly correlated with ataluren inhibition of RFC activity. As such, they provide an attractive hypothesis for the variability of ataluren effectiveness, suggesting that patients harboring a PTC mutation with a sequence context leading to very strong interaction with RFC (low K_M_^RFC^ or K_A_^RFC^ values) will likely be resistant to ataluren, whereas sequence contexts conferring weaker interaction with RFC (higher values of K_M_^RFC^ or K_A_RFC) would likely be more amenable to ataluren treatment. Such treatment could be with ataluren added alone, or combined with other nonsense suppressors such as ACE-tRNAs or small molecule organic drugs such as aminoglycosides. Although a recent clinical trial of ELX-02, a promising aminoglycoside TRID, gave disappointing results^**53**^, it is possible that ELX-02 would be more effective if used in combination with ataluren or other termination inhibitors.

For CF nonsense mutations, the possibility that the variability of patient response to ataluren is largely due to PTC sequence context effects of RFC activity could be tested by retrospective analysis of ataluren clinical trials in which the nonsense mutation within each patient’s CFTR mRNA was identified. Such information, which is not currently in the public domain, would indicate whether there is a correlation between the sequence context dependence of readthrough which we observe (Table 2, Figures 3, 5 - 7) and the response of individual CF patients to ataluren treatment. We appreciate the need to protect patient privacy in carrying out such an analysis, but we suggest that the effort required to achieve such protection would be justified, given the potential of a positive result which would enable pre-selecting CF patients most likely to benefit from ataluren treatment in future trials. The importance of directly linking PTC mRNA sequence context to treatment outcomes is of even greater importance for future trials which use ataluren as part of a combination therapy, and would likely benefit a larger number of CF patients. This number would increase still further if ongoing and future efforts are successful in discovering new TRIDs, which, like ataluren, would exclusively target termination activity at PTCs and retain ataluren’s low toxicity, but with lower EC_50_ values.

## ACKNOWLEDGEMENT

We thank the Christian Kaiser lab (Johns Hopkins University) for their gift of Sfp enzyme.

## FUNDING

This work was supported by research grants to B.S.C.(Cystic Fibrosis Foundation COOPER23G0; NIH RO1GM127374; NIH RO1HL160726) to Y.E.G ((R35GM118139) and to J.D.L RO1HL153988).

## SUPPLEMENTARY DATA

included below, consists of Figure S1 and Tables S1 – S2.

**Figure S1.**
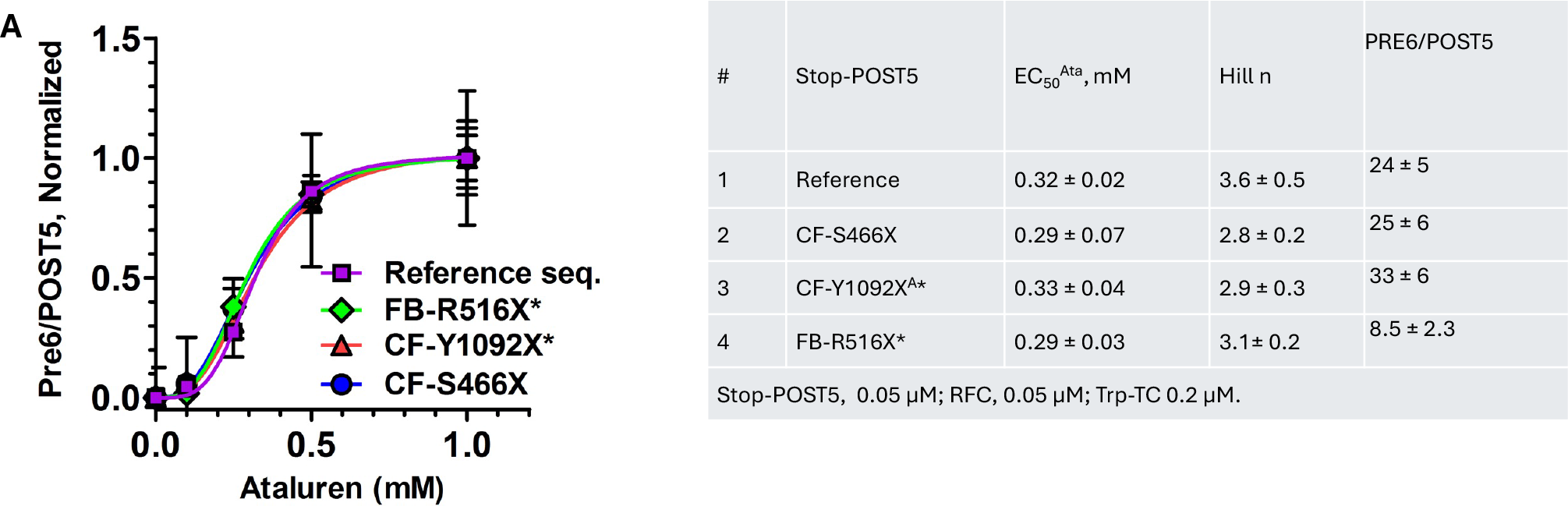
*A*. Similarity of dependence of readthrough on ataluren concentration for different mutant sequences. *B*. Parameter values.

**Table S1.**
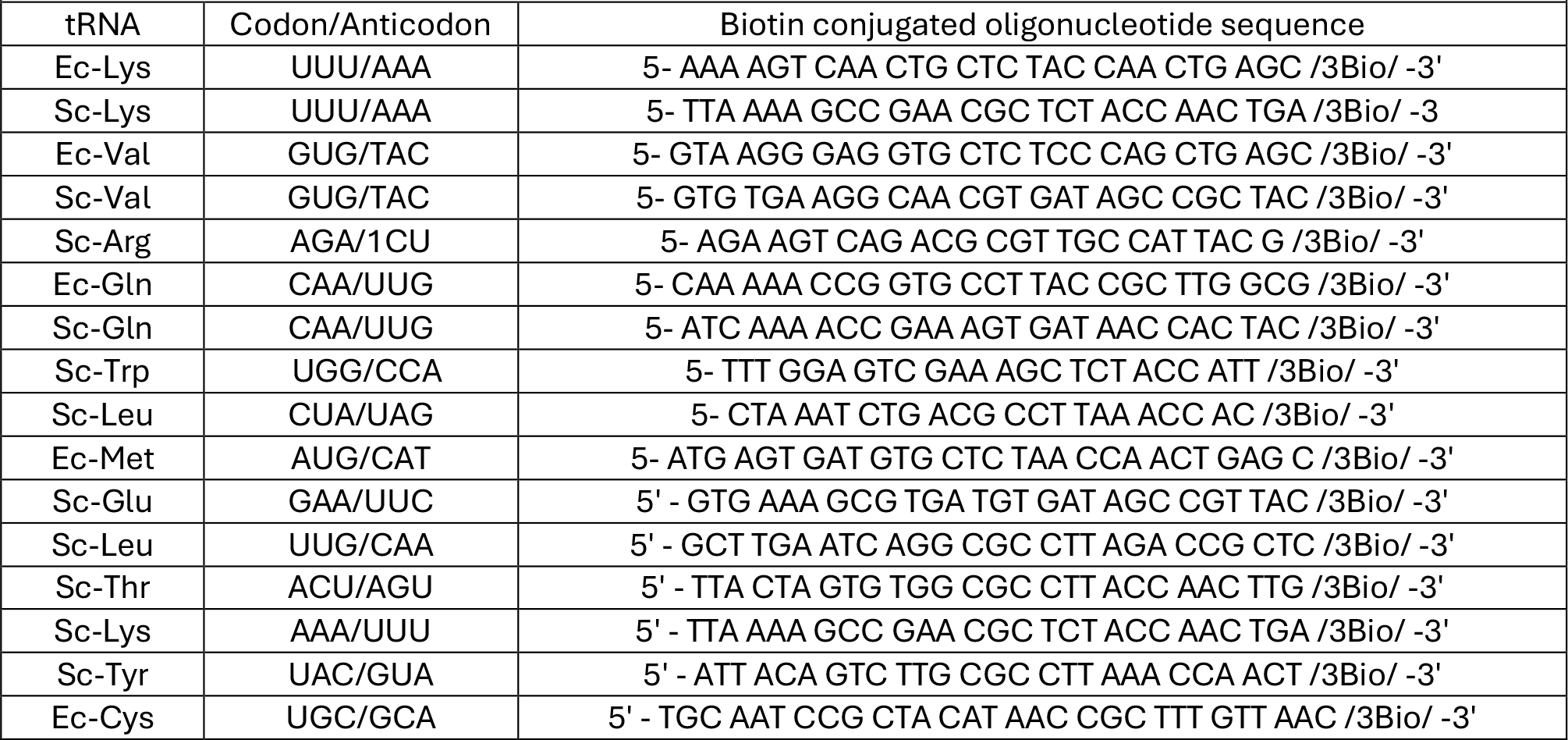
Oligo-DNA sequences used in purifying isoacceptor tRNAs.

**Table S2.** Rate constants for RFC interaction with Stop-POST5 complexes determined by smFRET experiments.

| Stop-POST5 | Sequence downstream | $k_{\text{arrival,app}}(\text{min}^{-1})$ | $k_{\text{heRF1 dis}}(\text{min}^{-1})$ | $k_{\text{tRNA dis}}(\text{min}^{-1})$ |
| --- | --- | --- | --- | --- |
| Reference | UGA CUA AUG | $1.63 \pm 0.14$ | $0.061 \pm 0.005$ | $0.065 \pm 0.005$ |
| CF-S1196X | UGA CAC GUG | $1.36 \pm 0.19$ | $0.049 \pm 0.006$ | $0.062 \pm 0.006$ |
| CF-W1282X | UGA AGG AAA | $1.50 \pm 0.15$ | $0.057 \pm 0.007$ | $0.065 \pm 0.006$ |
| CF-R1162X | UGA CUC UUU | $1.31 \pm 0.18$ | $0.039 \pm 0.006$ | $0.056 \pm 0.007$ |
| FB-R516X | UGA GCU GGA | $1.56 \pm 0.13$ | $0.043 \pm 0.005$ | $0.049 \pm 0.005$ |
| FB-R2694X | UGA GGA AAC | $1.23 \pm 0.17$ | $0.045 \pm 0.008$ | $0.059 \pm 0.008$ |
| CF-R553X* | UGA GCA AGA | $2.05 \pm 0.13$ | $0.014 \pm 0.003$ | $0.012 \pm 0.006$ |
| CF-RQ30X | UAG CGC CUG | $0.94 \pm 0.13$ | $0.034 \pm 0.006$ | $0.064 \pm 0.007$ |
| CF-Y1092X<sup>G</sup> | UAG CUG UCA | $1.41 \pm 0.12$ | $0.048 \pm 0.006$ | $0.055 \pm 0.007$ |
| CF-Q685X | UAA UCU UUU | $1.35 \pm 0.13$ | $0.051 \pm 0.006$ | $0.052 \pm 0.006$ |
| CF-E92X | UAA GUC ACC | $1.81 \pm 0.14$ | $0.038 \pm 0.004$ | $0.041 \pm 0.006$ |

## Notes

### Competing Interest Statement

The authors have declared no competing interest.

### Summary of Updates

Figures 5, 6, and 7 - minor changes Tables 1 and 2 - minor changes

